# Macrophage PIM1 Drives Atherosclerosis by Enhancing Foam Cell Formation Via CD36

**DOI:** 10.1101/2025.10.31.685966

**Authors:** Mirza Ahmar Beg, Quoc Quang Luu, Vaya Chen, Yaxin Wang, Gang Xin, Weiguo Cui, Roy L. Silverstein, Yiliang Chen

**Author notes:** These authors equally contributed to this work (Mirza Ahmar Beg, Quoc Quang Luu). **Corresponding Authors:** Yiliang Chen, PhD Medical College of Wisconsin, Milwaukee, WI, USA, 8701 West Watertown Plank Road Milwaukee, WI, 53226. Roy L Silverstein, MD Medical College of Wisconsin, Milwaukee, WI, USA, 8701 West Watertown Plank Road Milwaukee, WI, 53226. Weiguo Cui, PhD Northwestern University Feinberg School of Medicine, Department of Pathology, Chicago, IL, USA, 303 East Chicago Avenue, Chicago, IL, 60611 Total word count: 7259.

## Abstract

**Background:** Atherosclerosis is characterized by the buildup of fatty plaques that thicken and stiffen arterial walls. Macrophages (Mφs) significantly contribute to this process through their scavenger receptor CD36. PIM1 is a serine/threonine kinase known to modulate immune responses and cell metabolism. However, its role in Mφ lipid handling and atherogenesis is not well defined. This study examines the role of PIM1 in regulating CD36 expression and function in Mφs during foam cell formation and atherosclerosis progression.

**Methods:** We performed *in vitro* studies by treating murine peritoneal Mφs from *Pim1*^-/-^ and wild-type (WT) mice with oxidized low-density lipoprotein (oxLDL). We measured CD36, PIM1, and plaque-associated proteins and mRNA levels, oxLDL binding and uptake rates, as well as foam cell formation. For *in vivo* studies, we fed Mφ-specific PIM1-deficient (*Apoe*^-/-^ *Lyz2*^Cre/+^*Pim1*^fl/fl^) and their littermate control (*Apoe*^⁻/⁻^*Pim1*^fl/fl^) mice a high-fat diet for 12 weeks. We then evaluated the plaque formation in their aortic sinuses and arches.

**Results:** Deletion of *Pim1* in Mφs reduced CD36 protein expression by up to 96.7% compared to WT controls. This led to a 49.6% decrease in foam cell formation and a 25.5% reduction in cellular cholesterol after oxLDL treatment. Pharmacological inhibition of PIM kinase activity in WT Mφs also impaired oxLDL handling, with a 64.5% reduction in binding and a 57.9% in uptake. Bulk RNA-seq revealed that *Pim1* deficiency downregulated PPARγ signaling. Treatment with a PPARγ agonist restored CD36 levels in the PIM1 knockdown Mφs, suggesting that PIM1 regulates CD36 through PPARγ. Moreover, PIM1 Mφ-specific deficiency caused a 69.4% reduction in atherosclerotic plaque formation.

**Conclusion:** PIM1 acts as a key upstream regulator of CD36 by enhancing PPARγ activity in Mφs. The PIM1-CD36 axis promotes oxLDL binding, uptake, and foam cell formation. Targeting the PIM1/PPARγ/CD36 pathway could offer new ways to modulate Mφ lipid metabolism and reduce atherosclerotic plaque progression.

**Non-standard Abbreviations and Acronyms:** ELISA: enzyme-linked immunosorbent assay; HFD: high-fat diet; Mφs: macrophages; MCP-1: monocyte chemoattractant protein-1; ORO: oil red O; oxLDL: oxidized low-density lipoprotein; PBS: phosphate-buffered saline; PPARγ: peroxisome proliferator-activated receptor gamma; WT: wild type.

## Introduction

Atherosclerosis is a chronic inflammatory condition that impacts large and medium-sized arteries. It is characterized by the gradual buildup of immune cells, lipids, fibrous tissue, and calcium in the artery walls.^1,2^ Decades of intensive research have highlighted the central role of macrophages (Mφs) in the development of atherosclerosis. They perform essential and diverse functions throughout all stages of the disease, from initiation to plaque formation and rupture, by facilitating uptake of oxidized low-density lipoproteins (oxLDL), formation of foam cells, production of inflammatory substances, induction of oxidative stress, efferocytosis of dead and dying cells, and regulation of plaque stability.^3–5^ Previous studies, including ours, have shown that Mφs mediate production of proinflammatory cytokines in response to oxLDL through the type 2 scavenger receptor CD36.^6–8^ Recent preclinical animal studies highlight the therapeutic potential of targeting CD36 in atherosclerosis. Mφs-specific CD36 deficiency significantly reduced necrotic core area and improved plaque stability.^9–12^ Additionally, a major issue is that oxLDL/CD36 signaling initiates a positive feedback loop, increasing CD36 expression by activating the nuclear hormone receptor peroxisome proliferator-activated receptor gamma (PPARγ). This, in turn, boosts oxLDL uptake and ultimately leads to the formation of foam cells.^13^ However, the key regulator of this positive feedback loop remains unclear, making it challenging to develop targeted inhibitors for this pathway.

PIM1 is a constitutively active serine/threonine kinase that integrates inflammatory and metabolic signaling to influence cell differentiation, survival, and immune function.^14,15^ Although best known for its oncogenic roles, our previous work showed that PIM1 drives fatty acid oxidation and lipid uptake in myeloid-derived suppressor cells by sustaining PPARγ activity and expression of its downstream targets, including CD36.^16^ These findings prompted us to investigate whether PIM1 similarly controls lipid handling in arterial macrophages. Here, we hypothesize that PIM1 promotes macrophage foam-cell formation and atherogenesis by enhancing PPARγ-dependent CD36 expression and function.

Our results show that PIM1 is essential for both surface and intracellular CD36 expression in Mφs. In response to oxLDL, *Pim1*^-/-^ Mφs were resistant to foam cell formation and cholesterol accumulation. Using a new genetic mouse model (*Apoe*^-/-^*Lyz2*^Cre/+^*Pim1*^fl/fl^) and a high-fat diet (HFD) for 12 weeks, we demonstrated that *Pim1* Mφ-specific deficiency reduced plaque formation and pro-inflammatory cytokine MCP-1 in plasma. These findings support a pro-atherogenic role for PIM1 and underscore its potential as a therapeutic target against atherosclerosis.

## Materials and Methods

### Mouse Strains

All mice in this study had a C57BL/6 genetic background. Wild-type (WT) mice were obtained from Charles River Laboratories (Wilmington, MA, USA). *Pim1*^-/-^ (PIM1 full-body knockout) mice were originally obtained from Dr. Robert Woodland (University of Massachusetts). Germline deletion of *Pim1* in mice yields viable, fertile animals with only mild baseline phenotypes, such as erythrocyte microcytosis and subtle hematopoietic defects.^17^ These mice were later backcrossed to the C57BL/6 background for more than 12 generations. To interrogate macrophage-specific roles of PIM1 in atherosclerosis, *Pim1*^fl/fl^ mice were generated by GemPharmatech Co., Ltd (Nanjing, China) using CRISPR-Cas9-mediated genome editing. Founder mice were backcrossed to C57BL/6J mice to establish a stable *Pim1*^fl/fl^ line. *Apoe*^-/-^ and *Lyz2*^Cre/+^ mice were purchased from the Jackson Laboratory (Bar Harbor, ME, USA). To generate double-mutant mice, *Apoe*^-/-^ mice were crossed with *Pim1*^fl/fl^ mice. These double mutants were bred with *Lyz2*^Cre/+^ transgenic mice for several generations until Mφ-specific *Pim1*^-/-^ mice on an *Apoe*^-/-^ background (*Apoe*^-/-^*Lyz2*^Cre/+^*Pim1*^fl/fl^) were produced. Genotyping was conducted using PCR to confirm the presence of the floxed *Pim1* on both alleles, the *Cre* transgene, and the *Apoe* deletion. Littermate *Pim1*^fl/fl^*Apoe*^-/-^ mice were used as controls.

All animals were kept on a 12-hour light/dark cycle. They had free access to standard chow unless otherwise noted. For HFD experiments, adult mice of both sexes, aged 16–20 weeks, were randomly assigned to either continue on standard chow or switch to an atherogenic HFD (Harlan Teklad, Madison, WI, USA). This diet was provided for 6 or 12 weeks before analysis. The HFD contains 0.2% cholesterol and 21% total fat by weight, offering 42% of calories from fat.^18^ The control chow diet has only 5.1% total fat by weight and 13.6% of calories from fat. All procedures involving animals were performed in accordance with the guidelines and protocols approved by the Institutional Animal Care and Use Committee at the Medical College of Wisconsin.

### *In vivo* Foam Cell Assay

Peritoneal Mφs (8×10^6)^ from WT or *Pim1*^-/-^ mice were intraperitoneally injected into *Apoe*^-/-^ mice as previously described^19,20^ as a model of *in vivo* foam cell formation. The recipient mice had been fed an HFD for 6 weeks at the time of injection. In additional experiments, *Apoe*^-/-^ mice on the same HFD were transplanted with WT Mφs along with AZD1208 (MedChemExpress, MCE, Monmouth Junction, NJ, USA) at a dose of 30 mg/kg body weight by oral gavage. Control mice received only the vehicle. Peritoneal Mφs were collected three days after transfer, fixed with 4% paraformaldehyde at room temperature for 15 minutes^19^ and then stained with ORO for 5 minutes and imaged using a Motic system.

### Preparation of oxLDL

Copper oxidized human LDL was prepared as previously described.^18^ The degree of LDL oxidation was assessed using a thiobarbituric acid reactive substances assay kit (Abcam, Cambridge, UK). Protein concentration was measured by a bicinchoninic acid assay (Thermo Fisher Scientific, Waltham, MA, USA). oxLDL stock solutions were flushed with argon gas and stored at 4°C for further use. OxLDL was labeled with DiI as previously described.^21^ The labeled oxLDL was dialyzed to remove unbound DiI from the solution.

### Isolation And Treatment of Peritoneal Mφs

Mice received intraperitoneal injections of 2 ml of 4% thioglycolate (Sigma-Aldrich, St. Louis, MO, USA) in sterile saline, as previously described.^18^ Mφs were cultured in RPMI supplemented with 10% fetal bovine serum (FBS), 100 U/ml penicillin, and 100 μg/ml streptomycin, and maintained at 37°C in a humidified incubator with 5% CO₂. Cells were allowed to settle in complete media for at least two days before any experimental treatment, after which treatments were performed in RPMI with 0.2% MP Biomedicals™ Albumin, Bovine Fraction V, fatty acid free, low endotoxin, cell culture reagent (MP Biomedicals, LLC, Irvine, CA, USA). For experimental assays, Mφs from WT or *Pim1*^-/-^ mice were seeded into 6-well plates at 1 × 10⁶ cells per well.

#### Intracellular Staining, Surface Marker Analysis, oxLDL Binding and Uptake Assays Binding and uptake assays

To evaluate the effects of PIM inhibition on binding and uptake of oxLDL, isolated Mφs were treated with 10 μM AZD1208 for 72 hours. For the binding assay, the treated Mφs were incubated with 10 µg/mL DiI-oxLDL on ice for 30 minutes. For the uptake assay, the cells were incubated with 10 µM DiI-oxLDL at 37°C for 30 minutes. Surface-bound DiI-oxLDL was quenched with 0.2 M glycine (pH 2.0) and then neutralized with complete medium. Cells were washed twice with PBS to remove unbound particles, detached by scraping, and resuspended for flow cytometry analysis.

#### Intracellular and surface markers

Mφs were incubated with an anti-CD16 antibody for 30 minutes to block Fc receptors, followed by incubation with PE/Cyanine7 anti-mouse CD36 antibody (BioLegend, San Diego, CA) for 30 minutes to evaluate surface CD36 expression. For intracellular staining, cells were fixed and permeabilized using the eBioscience™ Foxp3/Transcription Factor Staining Buffer Set (Thermo Fisher Scientific) according to the manufacturer’s instructions, then stained with FITC-conjugated anti-PIM1 monoclonal antibody (Santa Cruz Biotechnology, Dallas, TX) for flow cytometry assays.

Flow cytometry was performed on a BD LSR II instrument using FACSDiva software with gain settings adjusted for each experiment based on unstained and single-color controls. Live cells were gated using forward and side scatter, and doublets were excluded via FSC-A versus FSC-H plots. Data were analyzed using FlowJo version 10.8.1 (Tree Star, OR).

### Aortic Lesion Analysis

For the aortic sinus and arch, a 0.24% ORO solution was prepared in 2-propanol and diluted with water. Dissected mouse aortic arches were cut open and fixed in 10% formalin overnight, then transferred to 1 mL of 10% formalin at 4°C for an additional overnight fixation. After washing, samples were *en face*-stained with ORO for 20 minutes, pinned, and examined under a stereomicroscope. Cardiac sinus tissues were immediately embedded in OCT compound (Fisher Healthcare, Pittsburgh, PA, USA) and sent to the Versiti Blood Research Institute Shared Resources Histology Core Facility for frozen section and slide preparation. After slide preparation, tissues were stained with ORO for 20 minutes, washed three times with PBS for 10 minutes each, and imaged using a Motic imaging system.

Cryostat sections were thawed at room temperature for 15 minutes and rehydrated in PBS for 10 minutes, then permeabilized with 0.2% Triton X-100 in PBS. Sections were subsequently incubated in blocking buffer (0.2% PBS-Triton X-100 containing 5% bovine serum albumin and 10% normal horse serum) for 1 hour at room temperature. Tissues were incubated overnight at 4°C in a humidified chamber with primary antibodies against mouse PPARγ (Santa Cruz Biotechnology), CD68 (Bio-Rad, Hercules, CA), or alpha-smooth muscle actin (αSMA, Thermo Fisher Scientific). After washing, sections were incubated with Alexa Fluor 488-conjugated goat anti-mouse or goat anti-rabbit antibody (Thermo Fisher Scientific) for 1 hour at room temperature, mounted with ProLong™ Diamond Antifade Mountant (Invitrogen), and imaged using a laser confocal fluorescence microscope (EVOS). Signals from each channel were quantified, and the total area of each color was measured using ImageJ. High-resolution images were captured using a Nikon widefield microscope.

### Transfection of siRNA

siRNAs targeting mouse *Pim1* were obtained from Integrated DNA Technologies (Coralville, IA) and incubated with INTERFERin® transfection reagent (Polyplus-transfection, Illkirch, France) for 10 minutes at room temperature. After a 2-day rest, Mφs were transfected with 50 nM siRNA duplexes in serum-free, antibiotic-free medium for 24 hours, followed by replacement with fresh complete medium for another 24 hours. Scrambled siRNA (Qiagen, Hilden, Germany) served as the negative control. In some experiments, siRNA-treated cells were pretreated with rosiglitazone (MedChemExpress) for 30 minutes before stimulation with oxLDL for 24 hours.

Mouse *Pim1* siRNA duplex sequence:

5′-rCrGrArGrArArCrArUrCrUrUrArArUrCrGrArCrCrUrGrAGC -3′

5′-rGrCrUrCrArGrGrUrCrGrArUrUrArArGrArUrGrUrUrCrUrCrGrUrC -3′

### RNA Extraction and Gene Expression Analysis

Cells were harvested for total RNA using the RNeasy^®^ Mini Kit (Qiagen, Hilden, Germany). cDNA was synthesized from the isolated RNA with the High-Capacity cDNA Reverse Transcription Kit (Applied Biosystems, Darmstadt, Germany). Quantitative PCR was conducted using PowerUp™ SYBR™ Green Master Mix (Applied Biosystems), with an annealing temperature of 60.6°C for all genes. Primer sequences are listed below:

### Mouse *SRA1*

5’-AACTCTGATAGAAGACGTGCTG-3’

5’-TGAGTGATCGGTGAATGTCATC-3’

### Mouse *SRB1*

5’-GTACCTCCCAGACATGCTTC-3’

5’-CTTGCTGAGTCCGTTCCATT-3’

### Measurement of Total Cholesterol and Inflammatory Cytokines

Plasma levels of total cholesterol and inflammatory cytokines were measured as previously reported.^21,22^

### Western Blotting

Proteins were extracted from Mφs using CelLytic™ M (Sigma-Aldrich) supplemented with cOmplete™, Mini, EDTA-free Protease Inhibitor Cocktail (Roche) and phosphatase inhibitor (Sigma-Aldrich) for 15 minutes. And then, they were incubated overnight at 4°C with primary antibodies against PIM1 (Cell Signaling Technology, Danvers, MA), CD36 (Novus Biologicals, Littleton, CO), SRA1 (Proteintech, Rosemont, IL), SRB1 (Novus Biologicals), and β-actin (Cell Signaling Technology) with gentle shaking.

### Bulk RNA-sequencing And Data Preprocessing

Total cellular RNA was isolated from peritoneal Mφs from WT or *Pim1*^-/-^ mice. RNA quality was evaluated with the Bioanalyzer RNA Nano Assay (Agilent). For RNA library preparation, the Illumina TruSeq Stranded mRNA kit (single indexed) was utilized, and sequencing was carried out on the Illumina HiSeq 2500 at the Versiti Blood Research Institute Core Facility. Each sample yielded an average of 40 million reads. Sequence reads were aligned with the STAR aligner 6 and the MAPR-Seq workflow v3.0. Differential expression analysis was performed using EdgeR v3.8 and the gene-wise exact test on TMM-normalized counts. Differential expression was determined by comparing each unique pair of conditions. Genes were deemed differentially expressed if the FDR-adjusted p-value was less than 0.05. Quality control was performed using FastQC and RSeQC 10, followed by manual review and data visualization.

### Reanalysis of Single-cell RNA-Sequencing Data

Single-cell RNA sequencing data from the GENE Expression Omnibus Database (GSE97310) were reanalyzed using the Seurat package.

### Statistical Analysis

The Shapiro-Wilk test was used to check for normal distribution. Data are presented as mean ± SD for normally distributed variables, while data are shown as median (25%–75%) for non-normally distributed variables. Differences between two continuous variables were evaluated using the Student’s t-test or the Mann-Whitney U test. For comparisons among multiple groups, the one-way or two-way ANOVA test was followed by Bonferroni’s post hoc test for normally distributed variables. Analyses were conducted with SPSS version 22.0 (IBM Corp., Armonk, NY, USA), and graphs were created with GraphPad Prism 6.0 (GraphPad Inc., San Diego, CA, USA).

## Results

### PIM1 is upregulated during atherosclerosis progression and facilitates CD36 expression in Mφs

Data from a previously published single-cell RNA sequencing study on HFD-fed *ldlr^-/-^* mice were analyzed to evaluate the association between *Pim1* expression in plaque Mφs and progression of atherosclerosis *in vivo*.^23^ Fig. 1A shows that *Pim1* expression was upregulated in the lesional Mφ clusters from both intermediate and advanced atherosclerotic aortas. OxLDL is a trigger of atherosclerosis because it primes inflammatory signaling and oxidative stress responses in Mφs, promotes their retention in plaque by inhibiting migration, and drives them toward foam cell formation, thereby promoting plaque progression and instability.^18,24–26^ CD36 is the primary receptor for Mφ oxLDL uptake,^3,12,27,28^ and treatment of Mφs with the PIM kinase inhibitor (AZD1208) resulted in a significant time-dependent decrease in CD36 expression. Compared to baseline, expression was decreased by ∼35% at 72 and 96 hours (*p* = 0.0007) (Fig. 1B). Consistent with the pharmacologic data, peritoneal Mφs isolated from *Pim1*^-/-^ mice showed a nearly total loss of CD36 surface expression (98.1% reduction compared to WT, *p* < 0.0001) (Fig. 1C). Total CD36 protein expression was also decreased by 94.2% in the *Pim1*^-/-^ cells as detected by immunoblot (Fig 1D, lane 3 compared to lane 1). Expression of other lipoprotein receptors (SRA1 and SRB1) was not changed in *Pim1*^-/-^ Mφs (Supplementary Fig. S1A-C).

**Figure 1.**
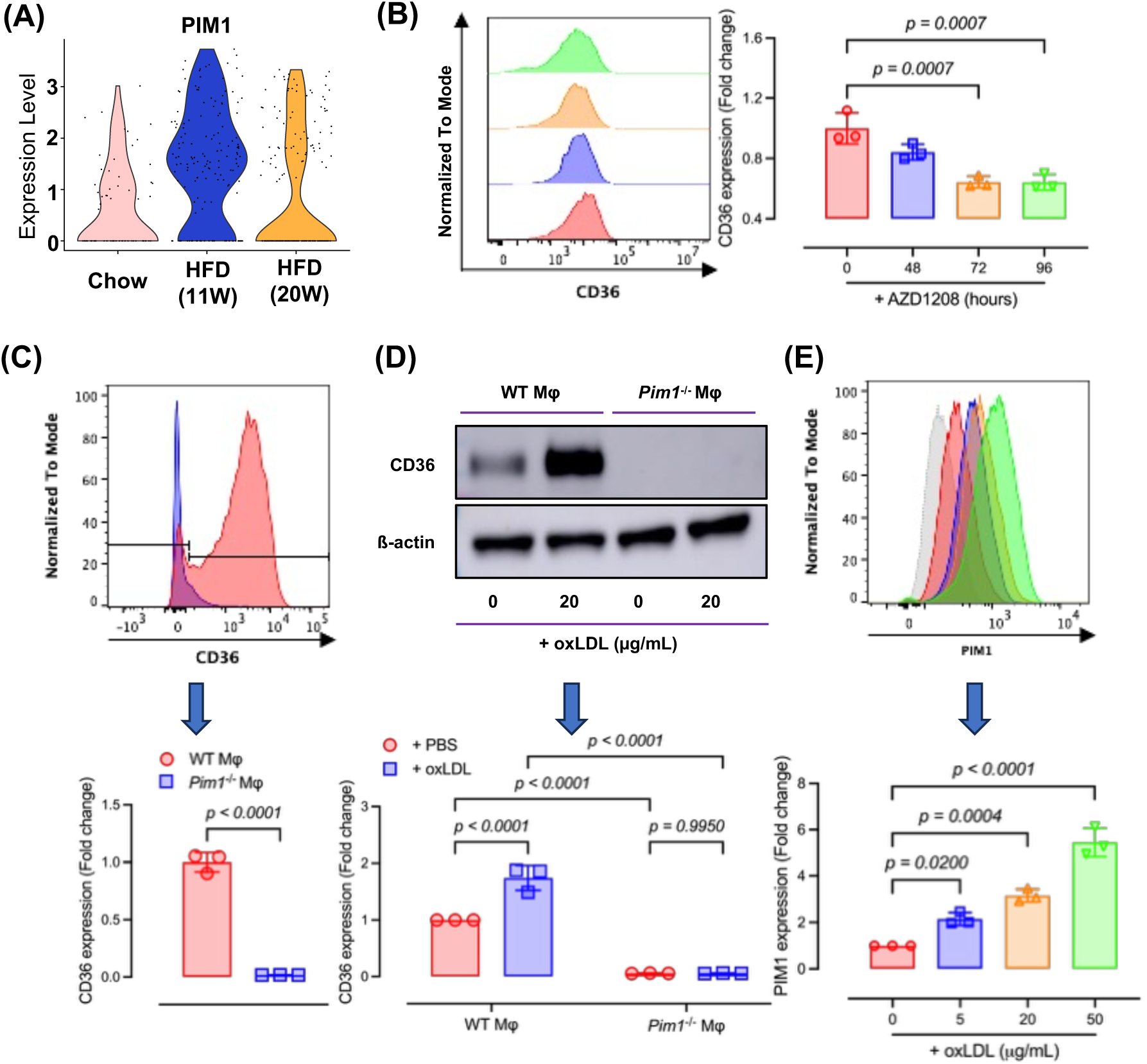
PIM1 is upregulated during the atherosclerosis progression and facilitates the expression of CD36 in Mφs. **(A)** Violin plots based on scRNA-seq show that *Pim1* mRNA was enriched and upregulated in plaque from *Ldlr*-null mice fed HFD. Each black dot represents an individual lesional Mφ. **(B)** Comparison of surface CD36 expression between vehicle-treated Mφs and AZD1208-treated Mφs. Examples of histograms of CD36 signals were shown. Flow cytometry data were quantified (n = 3 mice for each group). **(C)** Comparison of surface CD36 expression between peritoneal Mφs from *Pim1*^-/-^ mice and WT mice. Examples of histograms of CD36 signals were shown. Flow cytometry data were quantified (n = 3 mice for each group). **(D)** CD36 expression in WT and *Pim1*^-/-^ Mφ was evaluated by Western blot analysis. A representative blot image was shown on the left. Protein expressions were quantified by densitometry and shown in the bar graphs on the right (n = 3 mice for each group). **(E)** WT murine peritoneal Mφs were treated with increasing doses of oxLDL (0, 5, 20, and 50 µg/mL) for 24 hours and PIM1 protein expression was then evaluated using flow cytometry. Examples of histograms of PIM1 signals were shown. Mean fluorescence intensity (MFI) normalized to control is shown in the bar graph (n = 3 mice for each group).

CD36 is known to be up-regulated by oxLDL. As previously reported, we showed that oxLDL stimulation for 24 hours increased CD36 expression by 74.3% (*p* < 0.0001). This was not seen in *Pim1*^-/-^ cells (Fig. 1D, lane 4 compared to lanes 2 and 3). We also found that oxLDL stimulation of murine peritoneal Mφs *in vitro* significantly increased PIM1 protein expression in a dose-dependent manner as assessed by flow cytometry (Fig. 1E). At the highest dose tested (50 μg/ml), expression was increased by 5.45-fold (*p* < 0.0001). Collectively, these findings demonstrate that PIM1 is a key regulator of basal CD36 expression in Mφs and is an essential component of the oxLDL-mediated positive feedback loop that up-regulates CD36 expression and accelerates foam cell formation.

### PIM1 deficiency in Mφs reduces oxLDL binding and uptake, and inhibits foam cell formation *in vitro*

WT murine peritoneal Mφs pretreated with 10 µM of the PIM kinase inhibitor AZD1208 were incubated for 30min with DiI-oxLDL at either 4°C to assess surface binding or 37°C to assess uptake. Treatment with AZD1208 resulted in a reduction of both oxLDL-binding by 64.5% and uptake by 57.9% (Fig. 2A). We also assessed Mφ foam cell formation by ORO staining in WT and *Pim1*^-/-^ Mφs after 24-hour exposure to 50μg/ml oxLDL. While oxLDL stimulation nearly doubled foam cell formation in WT cells, this was reduced by 49.6% in *Pim1*^-/-^ cells (Fig. 2B-C; *p* < 0.005). Correspondingly, cellular cholesterol content measured by a biochemical assay was significantly reduced in *Pim1*^-/-^ Mφs (*p* < 0.005 for all; Fig. 2D). Together, these data suggest that PIM1 is required for CD36-mediated oxLDL binding, uptake and foam cell formation.

**Figure 2.**
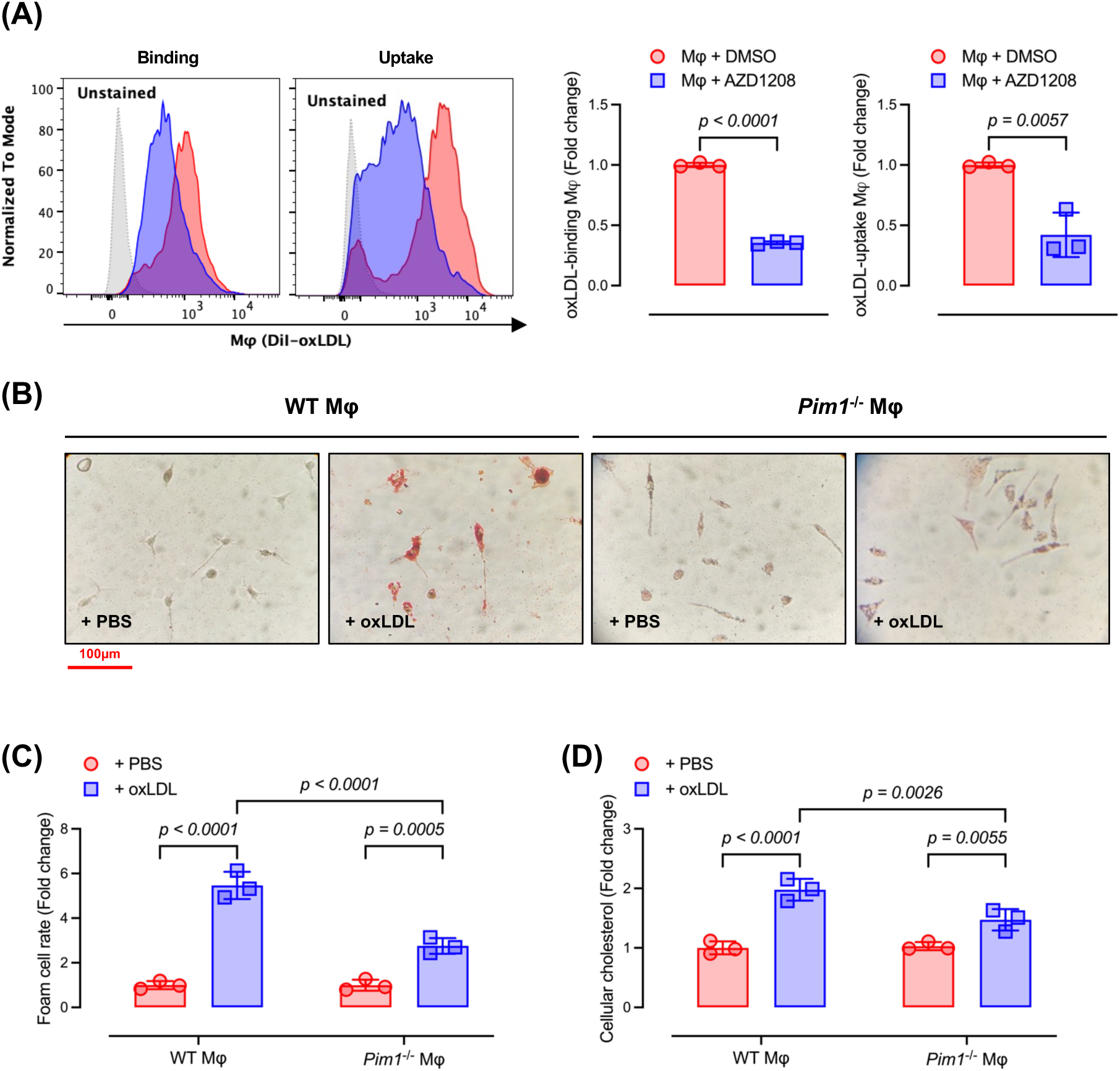
PIM1 deficiency in Mφs reduces oxLDL binding and uptake, which inhibits foam cell formation. WT peritoneal Mφ were treated with 10µM of AZD1208 or vehicle for 72 hours. **(A)** DiI-oxLDL-binding and -uptake assays by Mφ were evaluated by flow cytometry, respectively. Examples of histograms of DiI signals were shown. Flow cytometry data were quantified (n = 3 mice for each group). **(B)** Mφ from WT or *Pim1*^-/-^ cells were either treated with PBS or with 50µg/ml oxLDL for 24 hours. Oil Red O (ORO) staining was performed and representative images after staining were shown. **(C)** Foam cell rates were quantified (n = 3 mice for each group). More than 300 cells were counted for each condition. **(D)** Cholesterol was measured in peritoneal Mφs from WT and *Pim1*^-/-^ mice treated with PBS or oxLDL. Cholesterol concentrations were adjusted with protein content. Data in the bar graph were combined from three independent experiments.

### Macrophage PIM1 depletion or inhibition in hyperlipidemic mice attenuates foam cell formation *in vivo*

We used two independent approaches to provide *in vivo* validation of our in *vitro* experiments: intraperitoneal transplantation of *Pim1*^-/-^ Mφs and pharmacological inhibition of PIM1 with AZD1208 using the *Apoe*^⁻/⁻^ diet-induced hyperlipidemia murine model. As shown in Figure 3A-C, Mφs harvested from the intraperitoneal cavities of mice previously injected with *Pim1*^-/-^ Mφs exhibited a significant reduction in foam cell formation and intracellular cholesterol accumulation compared to Mφs harvested from mice receiving WT Mφs. In another experiment, mice injected with WT Mφs were treated with AZD1208 or vehicle via oral gavage for 3 consecutive days. The AZD1208-treated mice also showed decreased hyperlipidemia-driven foam cell formation and lipid deposition (Fig. 3D-F). Taken together, loss of PIM1 expression or activity in Mφs substantially reduces foam cell formation *in vivo*.

**Figure 3.**
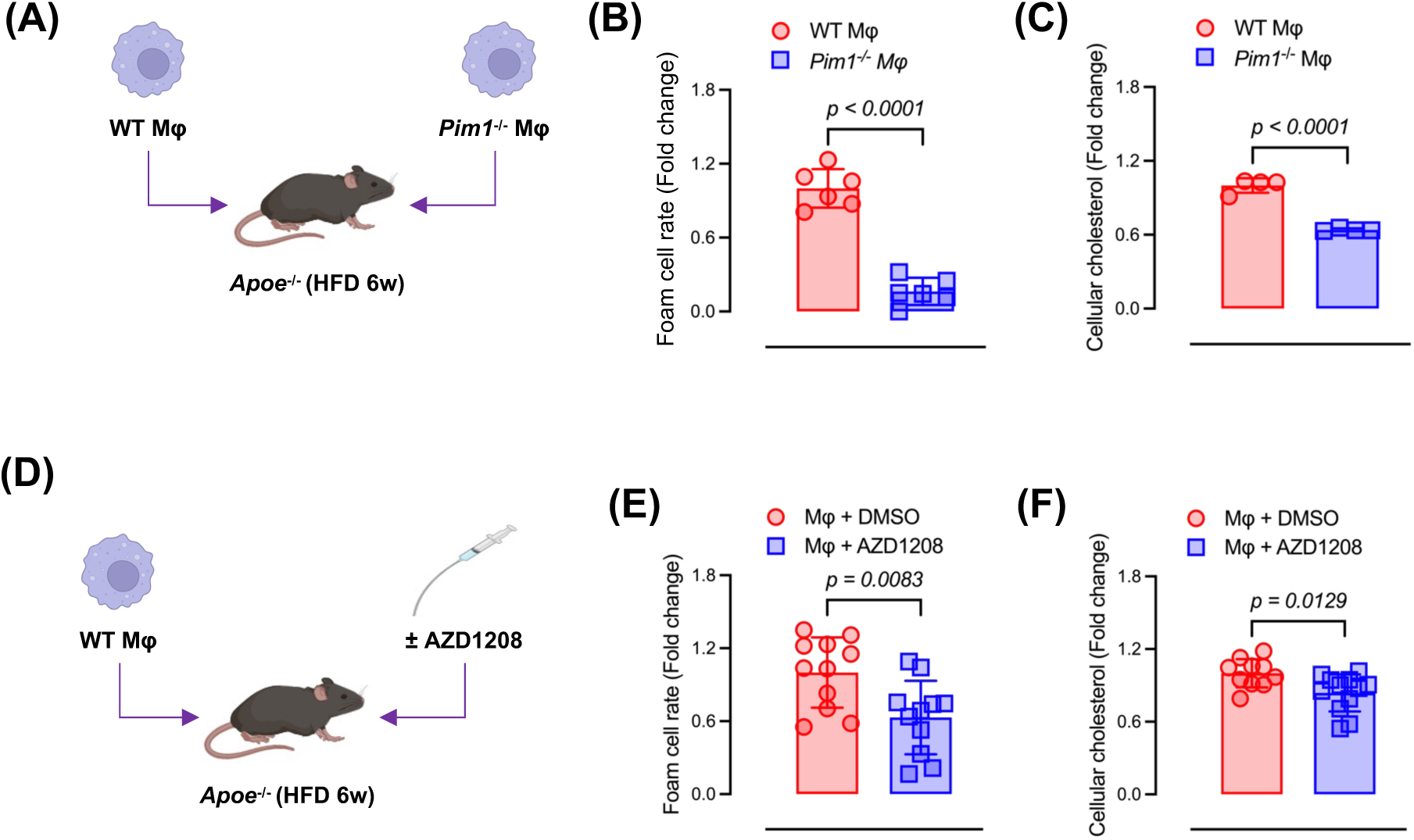
PIM1 deficiency in Mφs exhibits decreased foam cell formation *in vivo*. Mφ from WT or *Pim1*^-/-^ mice were injected intraperitoneally into *Apoe*^-/-^ mice fed on a HFD for 6 weeks to induce hyperlipidemia and a proatherogenic environment. **(A)** Schematic diagram showing the experimental design. **(B)** ORO staining was performed on peritoneal Mφs from HFD-treated *Apoe*^-/-^ mice transplanted with WT mice-or *Pim1*^-/-^ mice-derived Mφs. Foam cell formation was quantified using ORO staining (n = 6 mice for each group). More than 300 cells were counted for each condition. **(C)** Cholesterol was measured from cells in (B). Cholesterol concentrations were adjusted to protein content. Data in the bar graph were combined from four independent experiments. **(D)** *Apoe*^-/-^ mice on HFD for 6 weeks were injected with WT Mφ along with oral gavage of 30mg/g body weight of PIM inhibitor (AZD1208) or just vehicle. A schematic diagram showing the experimental design. **(E)** ORO staining was performed on peritoneal Mφs isolated from HFD-treated *Apoe*^-/-^ mice intraperitoneally injected with or without AZD1208. Foam cell formation was quantified by ORO staining (n = 11 mice for each group). More than 300 cells were counted for each condition. **(F)** Cholesterol was measured from cells in (E). Cholesterol concentrations were adjusted to protein content. Data in the bar graph were combined from 10-12 independent experiments.

### Mφ-specific PIM1 deficiency reduces atherosclerotic plaque formation in *Apoe*^-/-^ mice

To test whether PIM1 in Mφs contributes to atherosclerosis, we used a multi-step breeding strategy to generate Mφ-specific PIM1 deficient (*Apoe*^⁻/⁻^*Lyz2*^Cre⁺^*Pim1*^fl/fl^) and control (*Apoe*^⁻/⁻^*Pim1*^fl/fl^) mice. These mice were fed HFD for 12 weeks to induce hyperlipidemia and promote atherosclerotic lesion development (Fig. 4A). There was no significant difference between plasma cholesterol levels in the *Apoe*^⁻/⁻^*Lyz2*^Cre⁺^*Pim1*^fl/fl^ and *Apoe*^⁻/⁻^*Pim1*^fl/fl^ (*p* = 0.1088; Fig. 4B). Compared to *Apoe*^⁻/⁻^*Pim1*^fl/fl^ controls, the *Apoe*^⁻/⁻^*Lyz2*^Cre⁺^*Pim1*^fl/fl^ mice showed reductions in plasma MCP1 levels (*p* = 0.0243; Fig. 4C), consistent with a role for Mφ PIM1 in systemic inflammation. To assess plaque area, the aortic arches of the mice were dissected, opened *en face,* and stained with ORO. The *Apoe*^⁻/⁻^*Lyz2*^Cre⁺^*Pim1*^fl/fl^ mice displayed a 78.8% reduction in ORO-positive plaque area compared to *Apoe*^⁻/⁻^*Pim1*^fl/fl^ controls (Fig. 4D). Additionally, analysis of atherosclerotic plaque cross-sectional area and cellular content in sections obtained at the level of the aortic sinus revealed a 69.4% reduction in plaque area (*p* = 0.0185; Fig. 4E-F). Immunofluorescence-based assessments showed that CD68 expression decreased by 79.6% (*p* = 0.0330) (Fig. 4E and G), reflecting reduced Mφ content within the lesions. αSMA-fluorescence reflecting smooth muscle cell content was also ∼70% lower in *Apoe*^⁻/⁻^*Lyz2*^Cre⁺^*Pim1*^fl/fl^ mice compared to controls, but with a *p* value of 0.11 (Fig. 4E and G).

**Figure 4.**
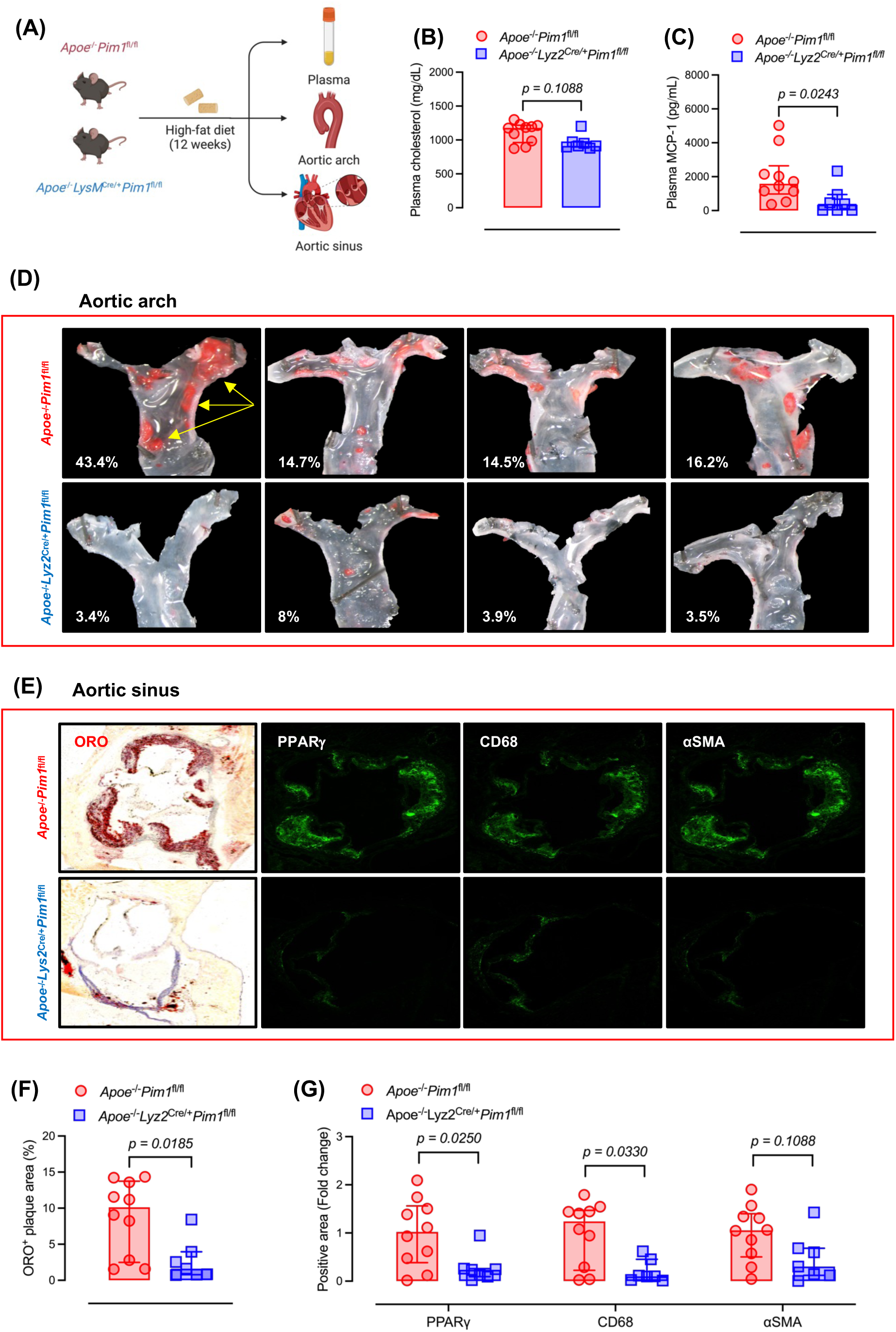
Mφ-specific *Pim1* deletion significantly reduces atherosclerotic plaque formation in *Apoe*^-/-^ mice. **(A)** A schematic diagram showing the experimental design. **(B)** The levels of plasma cholesterol (n = 7-10 mice for each group). **(C)** The levels of plasma MCP-1 (n = 7-10 mice for each group). **(D)** Aortic arches from four mice with *Apoe*^-/-^*Pim1*^fl/fl^ and *Apoe*^-/-^*Lyz2*^Cre/+^*Pim1*^fl/fl^ fed a HFD for 12 weeks were dissected, pinned open, and stained *en-face* for lipid-rich plaque with ORO. The yellow arrow indicates plaque. Representative images from four mice in each group are shown. Percent of plaque area was shown at bottom left corner in each image. **(E)** Left panels show ORO staining of histological sections obtained at the level of the aortic sinus from these mice with. Immunofluorescence images with antibodies to PPARψ, and CD68 are shown in the 3 panels to the right. **(F-G)** The quantifications of positive ORO, PPARψ, CD68, and αSMA areas in the aortic sinus were measured (n = 7-10 mice for each group).

Overall, these findings demonstrate that Mφ PIM1 is a key regulator of plaque formation and inflammatory responses during atherosclerosis.

### PIM1 modulates CD36 expression through PPARγ signaling

The PPARγ-CD36 axis is a key regulator in Mφ biology and atherosclerosis.^28,29^ Immunofluorescence staining with anti-PPARγ antibody in the aortic sinus sections showed that PPARγ expression was decreased by 72.3% (*p* = 0.0250) in *Apoe*^⁻/⁻^*Lyz2*^Cre⁺^*Pim1*^fl/fl^ compared to sections from the control mice (Figure 4G and E). Analysis of bulk RNA-sequencing data comparing WT Mφs to *Pim1*^-/-^ Mφs also showed reduced PPARγ expression, along with downregulation of its downstream targets CD36 and RXRα, which are essential for lipid uptake and nuclear receptor signaling (Fig. 5A and B). These data suggest that loss of PPARγ signaling could mediate the effect of PIM1 deletion on reducing CD36 expression.^28,29^ At the inflammatory axis, expression of transcripts for phosphoinositide-3-kinase (PI3K)/AKT serine/threonine kinase signaling intermediates (including *PIK3R1* and *AKT1*),^30^ proinflammatory regulators (*NFKB1*, *IKBKG*, and *TBK1*),^31^ and stress-response kinases (*MAPK3*, *MAP3K7*, and *MAPK9*),^32,33^ was also reduced in *Pim1*^-/-^ Mφs. This indicates that chronic PIM1 deficiency might blunt innate immune signaling, possibly affecting Mφ activation dynamics. On the metabolic level, *PPARGC1A*, a key regulator of mitochondrial biogenesis, was also decreased in *Pim1*^-/-^ Mφs (Fig. 5A).^34,35^ Moreover, enzymes involved in fatty acid oxidation (*ACADVL*, *ACADS*, *ACADSB*, *ACADM*, and *ACADL*) showed reduced expression, suggesting impaired mitochondrial lipid utilization in *Pim1*^-/-^ Mφs.^36^

**Figure 5.**
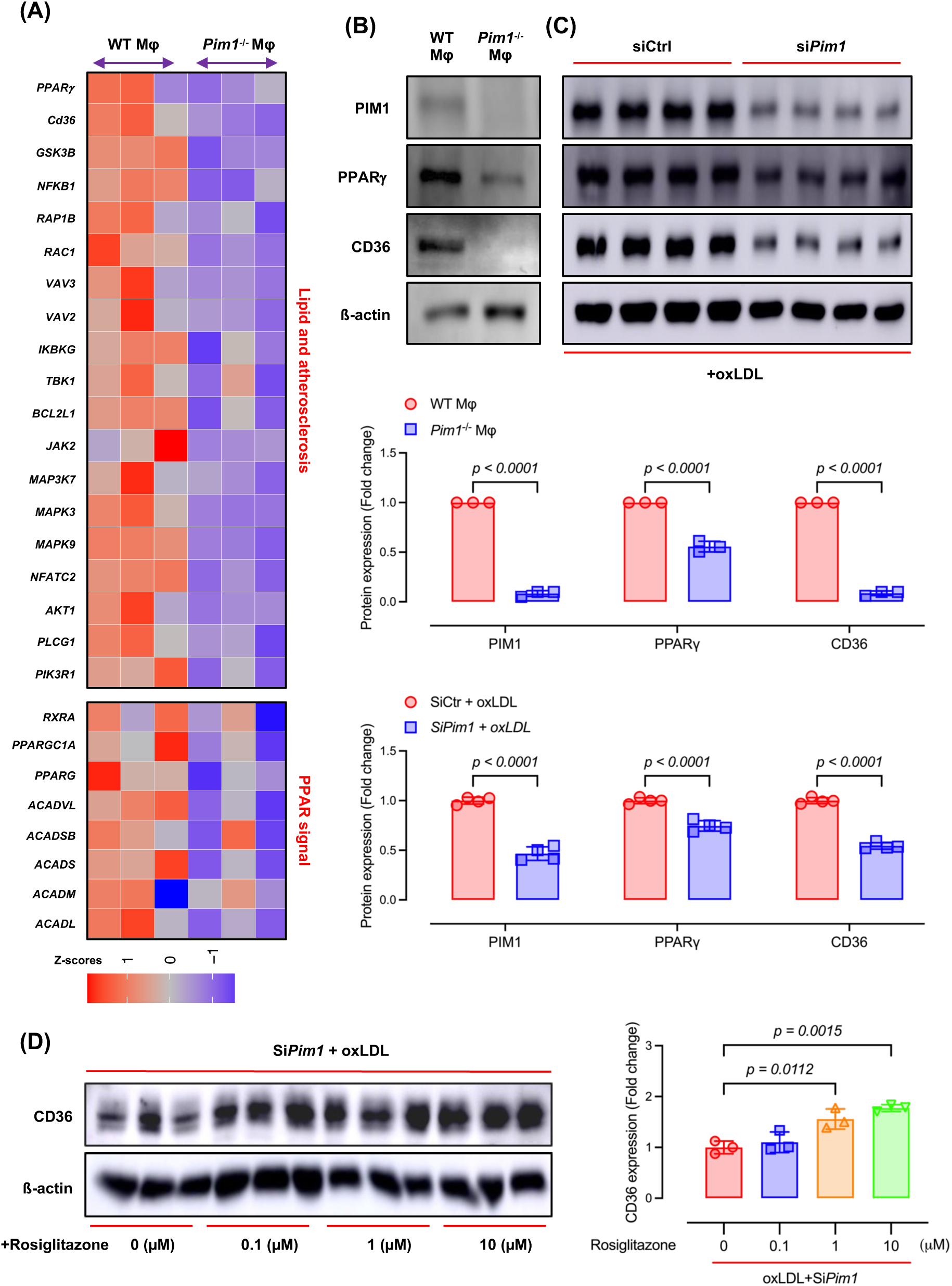
PIM1 modulates CD36 expression through PPARγ signaling. Peritoneal Mφs from WT and *Pim1*^-/-^ mice were isolated and cultured *in vitro*. Their RNA was then extracted and analyzed by bulk RNA sequencing. **(A)** A heatmap presents differentially expressed genes related to lipids, atherosclerosis, and PPAR signaling between WT and *Pim1*^-/-^ Mφ (n = 3 mice for each group). **(B)** Expression of PIM1, PPARγ, and CD36 in WT and *Pim1*^-/-^ Mφ were evaluated by Western blots. Representative blot images were shown (n = 3 mice for each group). **(C)** The effect of si*Pim1* on the expression of PIM1, PPARγ, and CD36 in Mφs was assessed by Western blots. Representative blot images were shown (n = 4 mice for each group). **(D)** The effect of rosiglitazone (a PPARγ agonist) on CD36 expression in Mφ was assessed by Western blots. Representative blot images were shown (n = 3 mice for each group).

To directly assess the role of PIM1 in regulating PPARγ and CD36, siRNA-mediated *Pim1* knockdown was performed in WT Mφs. To mimic an atherogenic condition *in vitro*, we pre-treated cells with oxLDL, followed by siRNA transfection. *Pim1* silencing significantly decreased both PPARγ and CD36 protein levels compared to control siRNA-transfected Mφs (Fig. 5C), indicating that oxLDL-stimulated PPARγ–CD36 axis requires PIM1. Conversely, treatment with a PPARγ agonist, Rosiglitazone, partially elevated CD36 expression in *Pim1*-knockdown Mφs (Fig. 5D). Overall, these findings suggest that PIM1 is upstream of the PPARγ/CD36 axis in Mφs in response to oxLDL.

## Discussion

Although dysregulated kinase signaling is increasingly recognized in cardiovascular disease, the roles of PIM1 in Mφ lipid metabolism and activation during atherosclerosis are poorly understood. This study provides *in vitro* and *in vivo* evidence that PIM1 is upregulated in Mφ under atherogenic conditions and is required for up-regulation of the scavenger receptor CD36. Mechanistically, PIM1 increases CD36 through the activation of PPARγ. This process promotes the pro-atherogenic Mφ phenotypes characterized by oxLDL binding, uptake, and foam cell formation, as well as pro-inflammatory signaling, thereby accelerating atherosclerotic plaque development.

Previous work has primarily linked PIM1 to cancer survival,^37^ and pro-inflammatory Mφ polarization.^38,39^ Our study demonstrates that PIM1 also drives Mφ lipid accumulation by maintaining the expression of CD36. Both genetic and pharmacological inhibition of PIM1 significantly reduced CD36 expression (Fig. 1). Notably, PIM1 deficiency nearly fully eliminated CD36 protein expression in Mφs, indicating the indispensable role of PIM1. Consistent with a predominant role of CD36 in oxLDL handling and lipid overloading,^25,40^ pharmacologic inhibition of PIM kinase activity with AZD1208 resulted in more than a 50% reduction in oxLDL binding and uptake (Fig. 2). Similar to CD36-deficient Mφs, PIM1-deficient Mφs were resistant to oxLDL-induced foam cell formation (Fig. 2 and 3).

CD36-deficient mice, either full body or Mφ-specific, are resistant to diet-induced atherosclerosis.^40,41^ To investigate the effects of specific *Pim1* deletion in Mφs during atherogenesis *in vivo*, we generated the *Apoe*^⁻/⁻^*Lyz2*^Cre⁺^*Pim1*^fl/fl^ mouse model. Deletion of Mφ *Pim1* resulted in decreased systemic inflammatory cytokines in plasma and reduced plaque formation in the aortic arch and sinus (Fig. 4). These findings are consistent with the notion that PIM1 promotes diet-induced atherosclerosis by facilitating CD36 expression in macrophages. Additionally, PPARγ activity and CD68⁺ Mφ burden were also reduced in the plaque from *Apoe*^⁻/⁻^*Lyz2*^Cre⁺^*Pim1*^fl/fl^ mice (Fig. 4). Circulating monocytes are recruited into the intima and differentiate into CD68⁺ Mφs.^42^ In these cells, PPARγ regulates lipid-handling pathways.^43,44^ The pathological outcome relies on the balance between lipid uptake, mainly mediated by CD36, and lipid disposal through cytoplasmic and mitochondrial mechanisms.^45^ Excessive lipid uptake results in cholesterol ester accumulation, foam cell formation, cholesterol crystal generation, and lysosomal stress, which activates NF-κB.^21,46,47^ This activation triggers the release of MCP-1, further recruiting monocytes. Collectively, the data show that the loss of *Pim1* disrupts the feed-forward loop involving PPARγ-driven lipid uptake, CD68⁺ Mφ accumulation, and NF-κB–mediated inflammation in the aortic sinus and arch. This disruption reduces foam cell formation and limits the progression of plaque.

Knockdown or knockout of *Pim1* in Mφs reduced the expression of PPARγ in the current study (Fig. 5) while pharmacological activation of PPARγ in *Pim1*-insufficient Mφs partially restored CD36 expression. Our results aligned with previous research showing that PIM1 phosphorylates STAT3 at Ser727, enhancing its transcriptional activity and increasing its binding to the PPARγ promoter. Inhibition of PIM1 with AZD1208 decreases STAT3 occupancy at PPARγ regulatory regions without significantly impacting other parts of the genome loci.^28,29,48,49^ Moreover, activated PPARγ forms heterodimers with RXRα and drives the expression of genes involved in lipid uptake, transport, and mitochondrial fatty acid oxidation.^34–36,44^ PPARGC1A functions as a transcriptional co-activator that encourages mitochondrial biogenesis and oxidative metabolism, while the acyl-CoA dehydrogenases (*ACADVL*, *ACADS*, *ACADSB*, *ACADM*, and *ACADL*) facilitate fatty acid breakdown of various chain lengths.^34–36,44^ Through this network, PPARγ couples lipid uptake to mitochondrial fatty acid oxidation, maintaining Mφ metabolic homeostasis through increasing CD36 expression. Nevertheless, dysregulation of this pathway by PIM1 deficiency may disrupt fatty acid breakdown and promote lipid-laden Mφ phenotypes.

Despite our new findings, several questions remain. First, it remains unclear how oxLDL upregulates PIM1 in Mφs. Previous studies have shown that the pro-inflammatory cytokine IL-6 induces PIM1 expression in pancreatic cancer cells.^50^ We reported that oxLDL/CD36 signaling activates NF-κB, leading to IL-6 production and secretion in Mφs.^22^ Thus, a plausible mechanism is that oxLDL/CD36-induced IL-6 acts in an autocrine manner through the IL-6 receptor to enhance PIM1 expression, which in turn sustains CD36 upregulation and establishes a positive feedback loop. Future studies are needed to test this hypothesis. Second, since CD36 protein level is almost fully eliminated in the PIM1-deficient Mφs, it is unclear whether PIM1 also regulates CD36 beyond the transcriptional level. The possibilities of PIM1 regulating CD36 protein stability or intracellular trafficking await future investigation. Third, it should be noted that AZD1208, used in our *in vitro* experiments to suppress PIM1 activity, is a pan-Pim kinase inhibitor that targets all three PIM isoforms (PIM1, PIM2, and PIM3).^51^ It limits our ability to ascribe its effects exclusively to PIM1, although the CD36 downregulation effect by AZD1208 is consistent with genetic PIM1 deficiency. In addition, the Lyz2^Cre^ system used for generating myeloid-specific PIM1-deficient mice is active not only in macrophages but also in neutrophils and other myeloid-derived cells. We cannot exclude the possibility that PIM1 deficiency in myeloid-derived cells other than macrophages partially contributes to plaque reduction in the *Apoe*^⁻/⁻^*Lyz2*^Cre⁺^*Pim1*^fl/fl^ mice.

In summary, our data demonstrate that PIM1 serves as a key regulator of foam cell formation and plaque progression, mediated by CD36-PPARγ signaling pathways. PIM1 inhibition might be a promising strategy to target Mφ-driven processes in atherosclerosis.

## Acknowledgments

The authors would like to acknowledge Versiti Blood Research Institute Shared Resources Core Facility (RRID: SCR_025503), supported by the Versiti Blood Research Institute Foundation, for their services, instrumentation, and specialist support. We especially thank our Histology Core staff Ms. Nicole Leon for the mouse tissue histology work. We also thank our animal technician, Mr. Douglas Franklin, for his excellent animal work.

## Data Availability Statement

All data generated for this study are included in this article. The raw data supporting the conclusions of this article will be made available by the authors on request.

## Author contributions

Q.Q.L. and Y. C. writing–original draft; Q.Q.L., R.L.S., W.C., and Y. C. writing–review and editing; Q.Q.L., M.A.B., V. C., Y. W., and Y. C. investigation; Q.Q.L., M.A.B., and Y. C. visualization; Q.Q.L. and M.A.B. methodology; R.S., W. C., and Y. C. funding acquisition; Q.Q.L., M.A.B., V. C., and Y. C. formal analysis; Q.Q.L., M.A.B., and V. C. data curation; Q.Q.L., R.L.S., and Y. C. conceptualization and experimental design; Y. C. project administration; Y. C. supervision; V. C. software. All authors have read and agreed to the published version of the manuscript.

